# Metabolic Alterations driven by PFKFB3 upregulation confer Resistance to Trastuzumab in HER2-Positive Breast Cancer

**DOI:** 10.1101/2025.10.09.681343

**Authors:** Roos Vincken, Samuel T. Pasco, Veronica Steri, Laura Bozal-Basterra, Wilfred F. J. van IJcken, Arkaitz Carracedo, Danny Huylebroeck, Mark M Moasser, Ana Ruiz-Sáenz

**Affiliations:** Department of Cell Biology, Erasmus University Medical Center Rotterdam, 3015 CN, Rotterdam, The Netherlands; Cancer Therapy Resistance Lab, Center for Cooperative Research in Biosciences (CIC bioGUNE), Basque Research and Technology Alliance (BRTA), Bizkaia Technology Park, Derio, Spain; Helen Diller Family Comprehensive Cancer Center, University of California, San Francisco, San Francisco, California; Preclinical Therapeutics Core, Helen Diller Family Comprehensive Cancer Center, University of California San Francisco, San Francisco, CA, USA; Cancer Cell Signaling and Metabolism Lab. CIC bioGUNE, BRTA, Bizkaia Technology Park, Derio, Spain; Center for Biomics, Erasmus University Medical Center Rotterdam, 3015 CN, Rotterdam, The Netherlands; Translational Prostate Cancer Research Lab, CIC bioGUNE-Basurto, Biobizkaia Health Research Institute.; Centro de Investigación Biomédica En Red de Cáncer (CIBERONC), Madrid, Spain; Biochemistry and Molecular Biology Department, University of the Basque Country (UPV/EHU), Bilbao, Spain; IKERBASQUE, Basque Foundation for Science, Bilbao, Spain

**Keywords:** HER2-targeting therapy, trastuzumab, ADCC, breast cancer, treatment resistance

## Abstract

**Aims:** Resistance to anti-HER2 therapies, particularly trastuzumab, remains a major obstacle in the treatment of HER2-positive (HER2+) breast cancer. This study aims to uncover novel mechanisms driving trastuzumab resistance with a focus on the immune component, key mediator of trastuzumab efficacy.

**Methods:** We developed an isogenic cell line-derived xenograft model to perform transcriptome-wide analyses of trastuzumab-sensitive and -resistant tumors. To validate key findings, we employed a 3D cancer–immune co-culture system capable of quantifying antibody-dependent cellular cytotoxicity (ADCC).

**Results:** Transcriptomic profiling revealed how trastuzumab treatment shifts tumor transcriptomes, including changes that remodel the metabolic landscape and distinct gene signatures associated with resistance, notably the upregulation of 6-phosphofructo-2-kinase/fructose-2,6-biphosphatase 3 (PFKFB3). Functional studies demonstrated that PFKFB3 promotes trastuzumab resistance by inducing metabolic rewiring and reducing ADCC. Silencing PFKFB3 restored immune-mediated cytotoxicity. Clinical dataset analyses confirmed that elevated PFKFB3 expression correlates with reduced overall and progression-free survival, and with incomplete pathological response to trastuzumab.

**Conclusions:** PFKFB3 upregulation drives metabolic adaptations that confer resistance to trastuzumab in HER2+ breast cancer. These findings highlight PFKFB3 as a promising therapeutic target to overcome resistance and improve patient outcomes.

**Highlights:**

- PFKFB3 upregulation promotes trastuzumab resistance in HER2+ breast cancer.
- Transcriptomic profiling reveals metabolic shifts linked to resistance.
- Silencing PFKFB3 restores ADCC and sensitizes cancer cells to trastuzumab.
- High PFKFB3 expression correlates with poor survival and treatment response.
- PFKFB3 is a potential therapeutic target to overcome trastuzumab resistance.

## Introduction

HER2-positive (HER2+) cancers are diagnosed in approximately one in five patients with breast cancer. These tumors are characterized by the overexpression of HER2, primarily caused by amplification of the *ErbB2* proto-oncogene, leading to constitutive signal transduction and eventually tumorigenesis. Activation of these oncogenic signaling pathways occurs through homo-or heterodimerization of HER2 with other receptor tyrosine kinases of the EGFR family of receptors, resulting in the recruitment of signaling proteins that drive cell proliferation and survival [1, 2].

Over the past decades, HER2-targeting agents, including antibodies, antibody-drug conjugates, and small-molecule tyrosine kinase inhibitors, have markedly improved the outcome of HER2+ breast cancer patients [3–5]. Trastuzumab, a humanized monoclonal antibody, has been the cornerstone of treatment for HER2+ cancers for more than three decades. Its therapeutic efficacy is attributed to its binding to the ectodomain of HER2 and its ability to engage the immune system, thereby promoting antibody-dependent cellular cytotoxicity (ADCC) [6].

The introduction of trastuzumab into clinical practice has significantly improved the survival rates. However, up to 42% of patients treated with neoadjuvant trastuzumab, and 27% of those treated with adjuvant trastuzumab, experience disease progression [7, 8]. To address this, extensive research has been conducted to uncover the mechanisms underlying trastuzumab resistance. These involve complex factors and pathways, including activating mutations in HER2 [9], heterodimerization with other growth factor receptors [10], truncated HER2 lacking the extracellular domain (ECD) [11, 12], loss of HER2 expression [13], and activation of alternative signaling pathways such as those involving estrogen receptors, c-Met, IGF1R (insulin-like growth factor-1 receptor) and AXL overexpression [14–17]. Downregulation of downstream signaling, especially in the PI3K pathway [18, 19], loss of PTEN expression [20], autocrine production of EGF-like ligands [21], and masking of the trastuzumab-binding epitope by MUC4 and hyaluronan [22–24] have also been implicated in resistance.

Despite these findings, overcoming trastuzumab resistance remains a challenge. To address this gap and identify novel clinically relevant mechanisms of trastuzumab resistance, we generated trastuzumab-resistant tumors using human cell line-derived xenografts in mouse models that retain immune cells critical to trastuzumab efficacy. Stimulation of the immune system is powerful in eliminating tumor cells, and trastuzumab engages the immune system, inducing ADCC. Our immunocompromised models include natural killer (NK) cells, which are critical mediators of ADCC. Through unbiased transcriptomic analyses of isogenic-resistant and-sensitive tumors, we discovered the upregulation of 6-phosphofructo-2-kinase/fructose-2,6-biphosphatase 3 (*PFKFB3)* in resistant tumors. Using our cancer-immune co-culture system, that is, combining cancer cells and NK cells [25], we found that silencing of *PFKFB3* increases NK cell-mediated ADCC. Next, we underscored the relevance of this finding through clinical data analyses showing that *PFKFB3* overexpression is inversely associated with pathological complete response (pCR), reduced overall survival (OS), and progression-free survival (PFS) in HER2+ breast cancer patients.

## Materials and Methods

### Animal studies

All animal experiments were performed by the UCSF Preclinical Therapeutics Core and were approved by the Institutional Animal Care and Use Committee at UCSF (IACUC). 6-to 7-female NCr nude mice (Taconic Biosciences, NCRNU-F (sp/sp), RRID:IMSR_TAC:NCRNU) were housed with ad libitum food and water on a 12-hour light cycle at the UCSF Preclinical Therapeutics Core vivarium.

### Trastuzumab resistant cell lines

To generate cell lines, the tumor was washed 3x with PBS and dissociated in DMEM:F10 without FBS using razor blades. The cell suspension was centrifuged at 200xg for 3 min and the supernatant was removed. Cells were resuspended in 1ml of 1mg/ml collagenase in 50mM TES 0.36 mM CaCl2 pH7.4 and incubated for 15 min at 37°C. Cells were diluted with 10 ml DMEM:F10 + 10% FBS + 1% PS, and single cells were selected by filtering through a 70 µm cell strainer. Cells were maintained at 37°C 5% CO_2_ and monitored for presence of fibroblasts.

### Trastuzumab resistant tumors

48 hours prior to tumor inoculation, E2 pellets (Innovative Research of America #SE1210.72MG_60_25P) were subcutaneously implanted in the scapular region. Subsequently, 4*10^6^ BT474 cells were injected subcutaneously and bilaterally in the dorsal flanks (serum free media and Matrigel (Corning #356237) mix 1:1); in NCr nude mice. The tumor dimensions and body weights were measured biweekly. Tumor volume was calculated to approximate the volume of an ellipsoid as 0.5 × width^2^ × length. Treatment enrollment was initiated on a rolling basis when the tumors reached 200mm^3^. Xenografted mice were randomly allocated to vehicle or trastuzumab arms and treated with vehicle (PBS) or 15 mg/kg trastuzumab (Herceptin) intraperitoneally twice a week. Resistant tumors were serially transplanted into treatment-naïve NCr nude female mice. Treatment with trastuzumab was resumed when the tumor reached a volume of 200 mm^3^.

### RNA isolation

RNA isolation for RNA sequencing was performed for six control tumors (C1,3-7), seven resistant tumors (T1-7), and two responders (T10 and T11). Four tumors were not RNA-sequenced because the mice died before a sample could be obtained or the tumor did not reach the lower cut-off of 100mm^3^ at the start of treatment. RNA isolation was performed using an RNeasy Plus kit (Qiagen #74134) according to the manufacturer’s instructions. For tumors 5-30 mg of tumor was homogenized in 600 μl RLT plus buffer in GenleMACS M tubes using the GentleMACS standard program RNA_02.

### RNAseq analysis

Sequencing libraries were created using the Truseq stranded mRNA library preparation method from Illumina and sequenced on an Illumina NextSeq2000 sequencer. The reads were aligned to the reference genome GRCh38 using HISAT2 (RRID:SCR_015530) [26]. After HISAT2 alignment, SAMTools (RRID:SCR_002105) [27] was used to sort and merge the alignments from multiple flow cells. Quantification was performed using HT-seq count [28] after first sorting the alignments on readnames using SAMTools. For mouse and human samples, only CCDS and mirbase transcripts were used for annotation. Non-CCDS genes and transcripts were removed, as many duplicate genes were present in the annotation, causing an increase in ambiguous alignments. These ambiguous alignments were not included in the expression profiles. The alignment percentage of the samples varied between 53-95% resulting in libraries with a minimum of 10M counts for each of the 13 samples. Two 2 samples (T11 and C7) had 5 M and 9 M counts in their profiles, respectively.

Count tables were read in RStudio (RRID:SCR_000432, version 2024.04.2), and genes with counts less than 10 were eliminated. The data were then processed using DESeq2 (RRID:SCR_015687, version 1.44.0) [29] for differential expression analysis. The list of identified differentially expressed genes (DEGs) was uploaded to DAVID (RRID:SCR_001881) for pathway analysis [30].

Gene set enrichment analysis (GSEA) [31] was performed by uploading normalized count tables into GSEA software (RRID:SCR_003199, version 4.3.3) using the Hallmarks (h.all.v2024.1.Hs.symbols.gmt) and Reactome (c2.cp.reactome.v2024.1.Hs.symbols.gmt) gene sets. All samples were compared using the Signal2Noise ranking metric; however, due to differences in sample size, comparisons with responding tumors used the Log2_Ratio_of_Classes ranking.

### RT-qPCR

RNA expression of PFKFB3 (Forward: TTGGCGTCCCCACAAAAGT, Reverse: AGTTGTAGGAGCTGTACTGCTT) was determined using RT-qPCR. cDNA was generated using iScript cDNA synthesis kit (BIO-RAD, #170-8890). qPCR was performed using 625 pg/μl cDNA, 300nM forward and reverse primers, and SYBR Green Supermix (BIO-RAD, #170-8886) in a 20μl reaction volume in triplicate. The reference genes GAPDH (Forward: GGTCTCCTCTGACTTCAACA, Reverse: AGCCAAATTCGTTGTCATAC) and ACTB (Forward: TGACGTGGACATCCGCAAAG, Reverse: CTGGAAGGTGGACAGCGAGG) were selected because of their stable expression across human breast cell lines [32]. Relative gene expression was compared using ordinary one-way ANOVA in GraphPad Prism (RRID:SCR_002798).

### Immunohistochemistry

Antigen retrieval was performed by heating to 100°C in 10mM Tris, 1mM EDTA buffer pH 9 with 0.05% Slides were blocked with 1% BSA in Tris-HCl buffer pH 7.6 and incubated with primary antibodies for detection of trastuzumab (Jackson ImmunoResearch Labs Cat# 109-035-098, RRID:AB_2337586, 1:500) or ki-67 (Agilent Cat# M7240, RRID:AB_2142367, 1:100) overnight at 4°C. Slides were incubated with secondary antibody for the detection of trastuzumab (Bethyl Cat# A50-100P, RRID:AB_67448, 1:100) and EnVision detection systems (Dako k5007) for ki-67 and developed with DAB (Dako). The slides were counterstained with hematoxylin and imaged using a Hamamatsu Nanozoomer 2.0 HT digital slide scanner.

The mean trastuzumab and ki-67 signal were quantified using ImageJ software version 1.54i (RRID:SCR_003070). Color deconvolution tool was used to separate the hematoxylin and DAB signals. The mean gray value in the DAB channel was calculated as the sum of the gray values of all pixels divided by the number of pixels. The mean trastuzumab signal was quantified in the whole slide after thresholding to remove the background signal. Mean ki-67 signal was quantified in 2-5 non-necrotic tumor regions of 250 nm2 for each tumor.

### Western blot

Tumor and cell lysates were prepared using modified RIPA lysis buffer (150 mmol/L NaCl, 0.1% SDS, 1% IGEPAL® CA-630, 1% sodium deoxycholate, and 10 mmol/L sodium phosphate, pH7.2) supplemented with protease inhibitors (Roche #05056489001) and phosphatase inhibitors (Roche #04906837001). Tumors C8, T8, and T9 were excluded because the mouse was found dead before a sample could be obtained.

Protein concentration was determined using BCA assay (Pierce #23228). 30 μg RIPA lysate was separated on an 8% SDS-PAGE gel and transferred to a PVDF membrane. Membranes were blocked with 3% BSA in 20 mM Tris, 136 mM NaCl buffer pH 7.4 (TBS), and incubated with primary antibodies diluted in 5% BSA in TBS with 0.05% Tween20 (TBS-T) overnight at 4°C. The membranes were washed with TBS-T and incubated with secondary antibodies for 1 h at room temperature. After washing with TBS-T, the antibodies were detected using fluorescence or chemiluminescence methods.

Western blotting was performed using antibodies purchased from Santa Cruz Biotechnology (HER2 Cat# sc-33684, RRID:AB_627996, HSP90 Cat# sc-13119, RRID:AB_675659), Cell Signaling Technology (p-HER2Y1248 Cat# 2247, RRID:AB_331725, pAKTThr308 Cat# 4056, RRID:AB_331163, pAKTSer473 Cat# 4058, RRID:AB_331168, p-MAPK Cat# 9101, RRID:AB_331646, MAPK Cat# 9102, RRID:AB_330744, GAPDH Cat# 2118, RRID:AB_561053, actin Cat# 3700, RRID:AB_2242334), and Abcam (p-PFKFB3 #202291, RRID:AB_3698749, PFKFB3 Cat# ab181861, RRID:AB_3095816). Horseradish peroxidase-conjugated secondary antibodies were purchased from Thermo Fisher Scientific (mouse Cat# G-21040, RRID:AB_2536527 and rabbit Cat# A16104, RRID:AB_2534776). Fluorophore-conjugated secondary antibodies were purchased from Li-COR Biosciences (mouse and rabbit, Cat# 926-68073, RRID:AB_10954442, Cat# 926-32212, RRID:AB_621847, Cat# 926-68072, RRID:AB_10953628, Cat# 926-32213, RRID:AB_621848). The signal was quantified using Image Studio Lite software (RRID:SCR_013715).

### Cell culture

All cells were maintained at 37°C and 5% CO2 and routinely tested for mycoplasma. BT474 (RRID:CVCL_0179), SK-BR-3 (RRID:CVCL_0033) and BT474-M1 derived cell lines were cultured in a 1:1 mixture of DMEM (Gibco #11965) and Ham’s F10 (Gibco #31550) +10% FBS (Capricorn Scientific #FBS-12A) +1% PS (Sigma #P0781). HCC1419 (RRID:CVCL_1251), HCC1569 (RRID:CVCL_1255), AU565 (RRID:CVCL_1074), HCC202 (RRID:CVCL_2062), HCC1954 (RRID:CVCL_1259), and UACC-893 (RRID:CVCL_1782) cells were cultured in RPMI 1640 (Gibco #52400) +10% FBS +1% PS. MDA-MB-361 (RRID:CVCL_0620) cells were cultured in DMEM +10% FBS +1% PS. SUM190PT (RRID:CVCL_3423) cells were cultured in DMEM:F10 +10% FBS +1% PS + 10μg/ml insulin + 1μg/ml hydrocortisone. KHYG-1 CD16+ cells were cultured in RPMI 1640 + 10% FBS + 1% PS + 2mM L-glutamine (Gibco #25030) + 1mM Sodium pyruvate (Gibco #11360) + 4.6 ng/ml IL-2 (PeproTech #200-02). Culture media for cell lines derived from trastuzumab-resistant tumors was supplemented with 0.15μg/ml trastuzumab. Assessment of HER2 dependency was performed by treating cells with 1μl lapatinib (LKT Labs #231277-92-2) or DMSO control for 2 h.

### Cell-viability assays

The antibodies trastuzumab (Herzuma/Trazimera) or IgG (Thermo Fisher Scientific Cat# 31154, RRID:AB_243591) were added in 10 dilutions (from 0.01-1000μg/ml) and two technical replicates in 96 well plates. The cells were seeded at a density of 3000 cells/well in culture medium. Cells were treated with antibodies in DMEM:F10 +1% FCS +1% PS for six days. Each cell line was screened for at least three biological replicates. Cell viability was measured using the CellTiter-GLo assay (Promega Corporation, #G7571), where the luminescence signal was read using a plate reader. Cell viability was converted into relative response using the average of untreated cells as control. Drug responses were plotted in GraphPad Prism version 10.1.2 (RRID:SCR_002798), and nonlinear regression analysis was used to fit a curve and obtain half-maximal inhibitory concentration (IC_50_) values. The model used is log(inhibitor) vs. response – variable slope (four parameters) with the bottom constrained to ‘shared value for all data sets’ and the top constrained to 100.

### Transfections

siRNA transfection was performed using Lipofectamine 2000 (Invitrogen #11668-019) according to the manufacturer’s instructions (Invitrogen). Control siRNAs were obtained from IDT (#51-01-14-04), and siRNAs targeting PFKFB3 were obtained from Dharmacon (#J-006763-05, #J-006763-07). ADCC assays were performed 72 h after knockdown.

### ADCC assay

Trastuzumab sensitivity was characterized as previously described [25]. Briefly, 1000 cells per well were seeded in a 96 well plate with or without 1 µg/ml trastuzumab. Tumor cells were incubated with KHYG-1 CD16+ cells at a 1:5 ratio for 6 h or a 1:3 ratio for 24 h at 37°C. Cell killing was measured using the Cytotox96 non-radioactive cytotoxicity assay (Promega #G1780) and cytotoxicity was calculated using the following formula: % cytotoxicity = (Experimental lysis - effector spontaneous - target spontaneous) / (target maximum - target spontaneous) × 100.

### Seahorse Glyco Stress metabolism measurements

Cells were plated in 24-well Seahorse XF cell culture plates (Seahorse Bioscience, 100882-004). Seahorse XFe24 sensor cartridge plates were hydrated with the XF Calibrant (Seahorse Bioscience) the day before the analysis and incubated overnight at 37°C without CO2. Before the bioenergetic measurements, the cells were washed and incubated for 1 h with Glyco Stress medium. Glyco Stress media contained XF Base Medium (minimal DMEM, Seahorse Bioscience, 103575-100) supplemented with 2 mM L-glutamine (Seahorse Bioscience, 103579-100). The extracellular acidification rate (ECAR), representative of glycolytic capacity, and the oxygen consumption rate (OCR), representative of mitochondrial respiration, were determined using the XFe Extracellular Flux Analyzer (Agilent/Seahorse Bioscience). The glycolytic metabolism was determined by sequential injection of 10 mM D-(+)-glucose (Sigma-Aldrich, G8644), 1.5 µM of the ATP synthase inhibitor oligomycin (Sigma, 75351) to inhibit mitochondrial respiration and force the cells to maximize their glycolytic capacity, and 50 mM 2-deoxy-D-glucose (2-DG) (Sigma-Aldrich, D8375), a competitive inhibitor of the first step of glycolysis. The concentrations indicated for each injection represent the final concentrations in the wells. At least three measurement cycles (3 min of mixing and 3 min of measurement) were performed before and after each injection. The OCR and ECAR were calculated using Wave software v2.6.3 (Agilent/Seahorse Bioscience). Following the manufacturer’s instructions, energy metabolism was normalized to the cell number using crystal violet staining.

### Clinical Data and Tissue Database Analysis

Kaplan-Meier plots were generated using the publicly available KM Plotter online tool (https://kmplot.com/analysis/), using breast cancer RNA sequencing and gene chip datasets [33].

Tissue expression was analyzed using the online tool from the Adult Gene Tissue Expression (GTEx) database and visualized using the data available for download (https://www.gtexportal.org/home/) [34].

Fully normalized and batch-corrected count tables from the ISPY2 clinical trial [35] were obtained from the Gene Expression Omnibus (GEO) database (RRID:SCR_005012) from SubSeries GSE194040, and mean expression values between HER2+ patients achieving pathological complete response (pCR) were compared using the Wilcoxon test. Transcriptional data of baseline and post-treatment samples from patients receiving trastuzumab were obtained from the GSE76360 [36], GSE114082 [37], and GSE245132 [38]. The mean expression values of samples between time points from GSE76360 and GSE114082 were analyzed using the Wilcoxon test. Differential expression analysis of GSE245132 was performed using the limma package (RRID:SCR_010943).

### Statistics

Statistical analyses were performed using GraphPad Prism Software (RRID:SCR_002798). Data are presented as mean ± SEM values of three or more biological replicates. Statistical significance was determined using a two-tailed unpaired test for immunohistochemistry and western blot quantification. For cytotoxicity experiments, statistical significance was determined using a two-tailed ratio paired t-test. Significant P-values are indicated with asterisks, ∗, P < 0.05; ∗∗, P < 0.005; ∗∗∗, P < 0.0005. Nonsignificant P values are marked as ns.

### Data availability

The transcriptomic data generated in this study (raw count table) are publicly available in the Mendeley Data repository at https://data.mendeley.com/datasets/nnp5h2pt8b/1.

## Results

### Generation and characterization of trastuzumab-resistant cells generated *in vivo*

To identify vulnerabilities that would aid in overcoming therapeutic resistance, we established a preclinical HER2+ breast cancer model of trastuzumab resistance. To this end, we generated trastuzumab-resistant tumors using a cell line-derived xenograft (CDX) model based on the prototypical HER2+ breast cancer cell line BT474, derived from a female invasive ductal breast carcinoma. In brief, BT474-M1 cells, derived from a passage of parental BT474 cells in mice, were implanted subcutaneously into nude female mice (Fig. 1A). Xenografted mice were treated with trastuzumab (15 mg/kg, twice a week) or vehicle control (Fig. 1A; C tumors). Among the trastuzumab-treated tumors (T tumors), three (#T1, T3, and T4) were unresponsive to therapy, whereas eight initially responded (Fig. 1A,B). Consistent with previous studies [39, 40], tumor regrowth was observed around day 35 in six of the eight initially responsive tumors despite continuous trastuzumab therapy. Tumors from these mice were surgically excised at necropsy for further analysis to investigate the mechanisms underlying their regained tumorigenic properties. Intrinsic resistance was not due to trastuzumab delivery failure, as resistant tumors showed evidence of trastuzumab within the tissue (Suppl. Fig. 1A-C). Resistant tumors also showed Ki67-positive proliferating cells, comparable to those in control tumors (Suppl. Fig. 1D-F). Furthermore, tumor regrowth was not driven by hyperactivation of the PI3K or ERK pathways, which are established mechanisms of resistance [18–20], as confirmed by western blot analysis of the tumor lysates (Fig. 1C,D).

**Figure 1.**
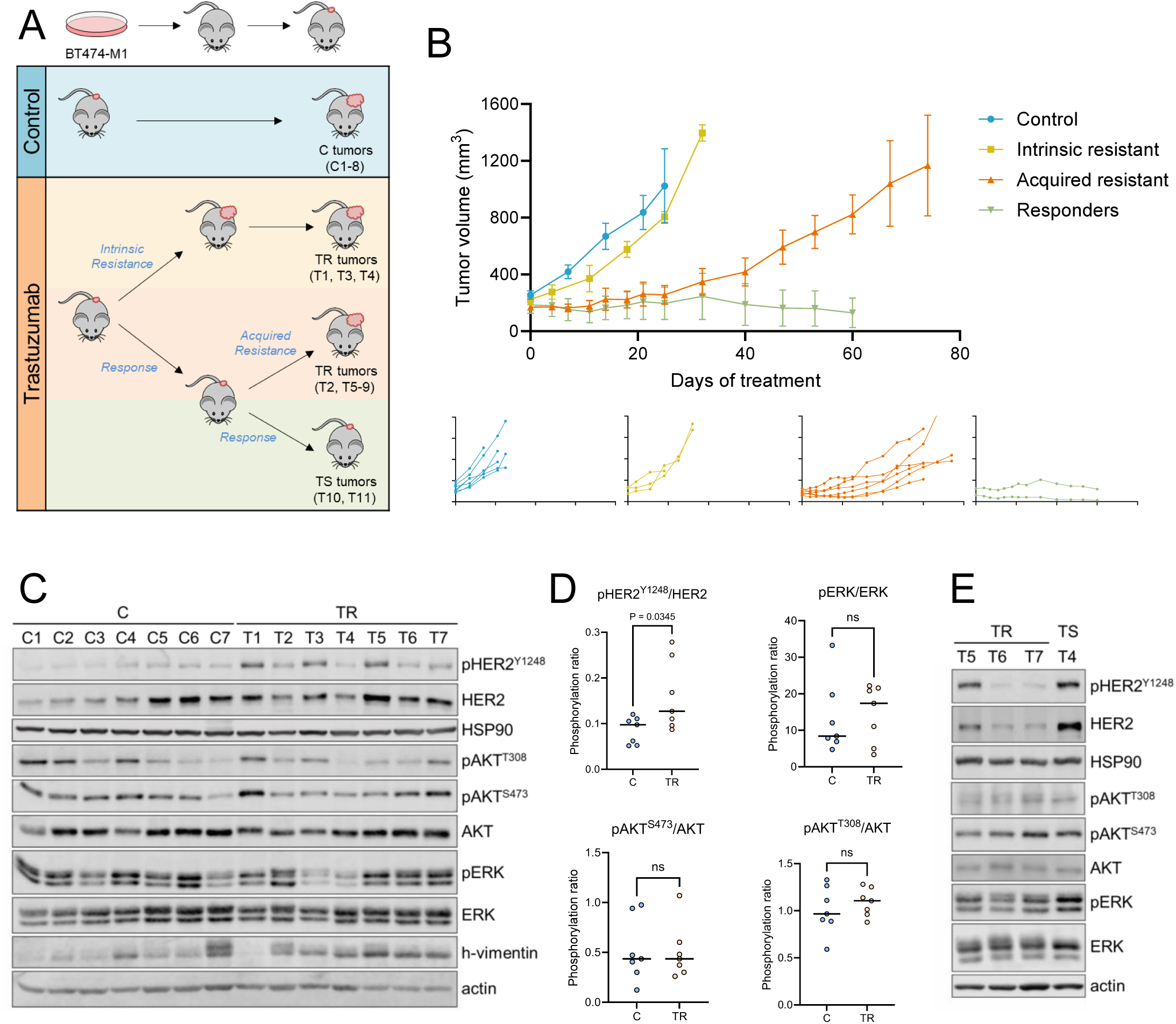

Interestingly, a tumor that remained sensitive to trastuzumab throughout the study exhibited PI3K and ERK pathway activation at levels comparable to those of resistant tumors (Fig. 1E; Suppl. Fig. 1G). Trastuzumab is known to exert its anti-tumor effects both by inhibiting these signaling pathways [20, 41] and by inducing antibody-mediated cellular cytotoxicity (ADCC) [6]. Thus, in the tumor that remains sensitive, cell intrinsic mechanisms are unlikely to be mediating the response. Instead, cell-extrinsic mechanisms—such as ADCC—are more likely to be involved.

### Transcriptomic profiling of trastuzumab-treated versus naive tumors

To explore the transcriptional differences between xenograft tumors, RNA was extracted from trastuzumab-treated and naïve tumors, and processed using RNA sequencing (RNAseq). Samples generally clustered according to treatment and outcome, with some variance (Fig. 2A). First, we compared trastuzumab-treated tumors with control tumors by differential gene expression analysis using a false discovery rate (FDR) < 0.05, to detect all significant differences. The treated tumors had 459 differentially expressed genes (DEGs): 280 upregulated and 179 downregulated (Fig. 2B). Gene set enrichment analysis (GSEA) of the dataset using the “Hallmarks” databases demonstrated that treated tumors were enriched for “Hypoxia” and metabolic processes (including “Oxidative Phosphorylation” and “Glycolysis”) (Fig. 2C). When comparing treated group subsets (responders, intrinsic resistance, and acquired resistance) to controls to find common signatures of treatment with trastuzumab, we found 29 upregulated and 14 downregulated genes (Fig. 2D). Interestingly, acquired resistant tumors had the fewest transcriptional differences compared to untreated tumors. These results demonstrate that trastuzumab treatment induces detectable shifts in tumor transcriptomes, including changes in metabolism.

**Figure 2.**
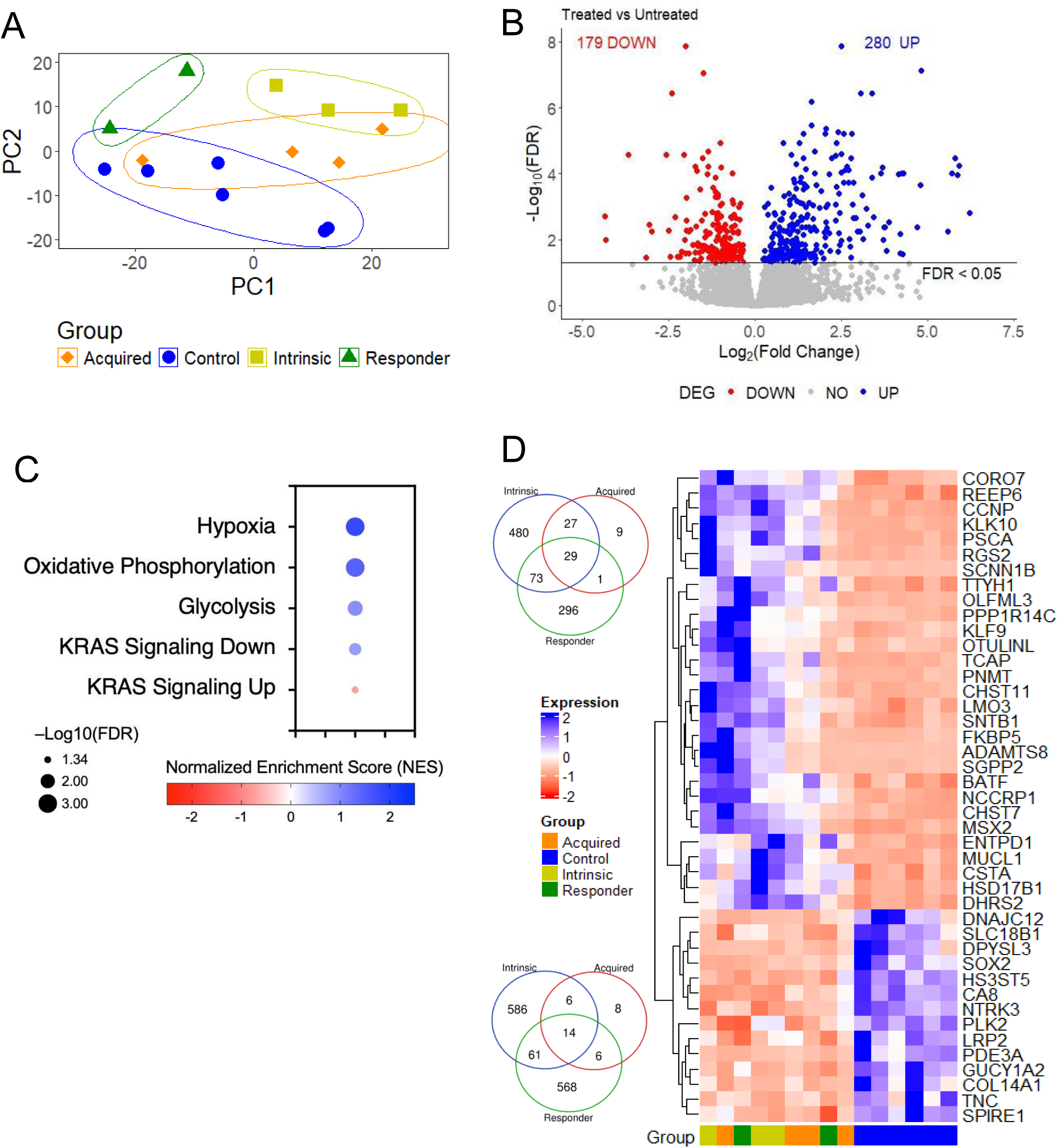

### Identification of molecular mechanisms of trastuzumab resistance

Next, we analyzed the transcriptomic profiles of tumors that received trastuzumab. This analysis revealed cell signaling differences between resistant and responsive tumors, providing potential hits involved in the development of resistance over time through comparative analysis of isogenic cancer xenografts. When comparing all resistant tumors to responding tumors, we adjusted the cutoff to include log fold change (LFC) thresholds of ±1.5 and found 500 upregulated and 43 downregulated DEGs (Fig. 3A). GSEA of this dataset using both the “Hallmarks” and “Reactome” databases revealed resistant tumors enriched for processes related to glycosylation, cytokine and chemokine signaling, and cell division whereas responder tumors were enriched for oxidative phosphorylation and starvation (Fig. 3B). When comparing responders to the resistant group subsets (intrinsic or acquired) to find common signatures of trastuzumab outcome, we found 252 upregulated and four downregulated genes shared (Fig. 3C). The expression of key genes, including those with the highest significance and fold change in the resistant group subsets and responding tumors are shown in Fig. 3D. Pathway analysis of these commonly upregulated DEGs revealed an overrepresentation of key processes for cellular growth in the resistant tumors (Fig. 3E). Among the genes significantly associated with resistance, we focused on *PFKFB3* upregulation (LFC = 2.46, FDR = 9.55e^-08^) as a potential resistance mechanism. Expression of *PFKFB3* in the breast was the fifth highest (out of 30 tissues) according to the GTEx Database [34] (Fig. 3F). *PFKFB3* encodes a glycolytic enzyme that converts fructose-6-phosphate to fructose-2,6-bisphosphate and, we confirmed elevated *PFKFB3* expression in both intrinsic and acquired resistant tumors by qPCR of tumor RNA (Fig. 3G).

**Figure 3.**
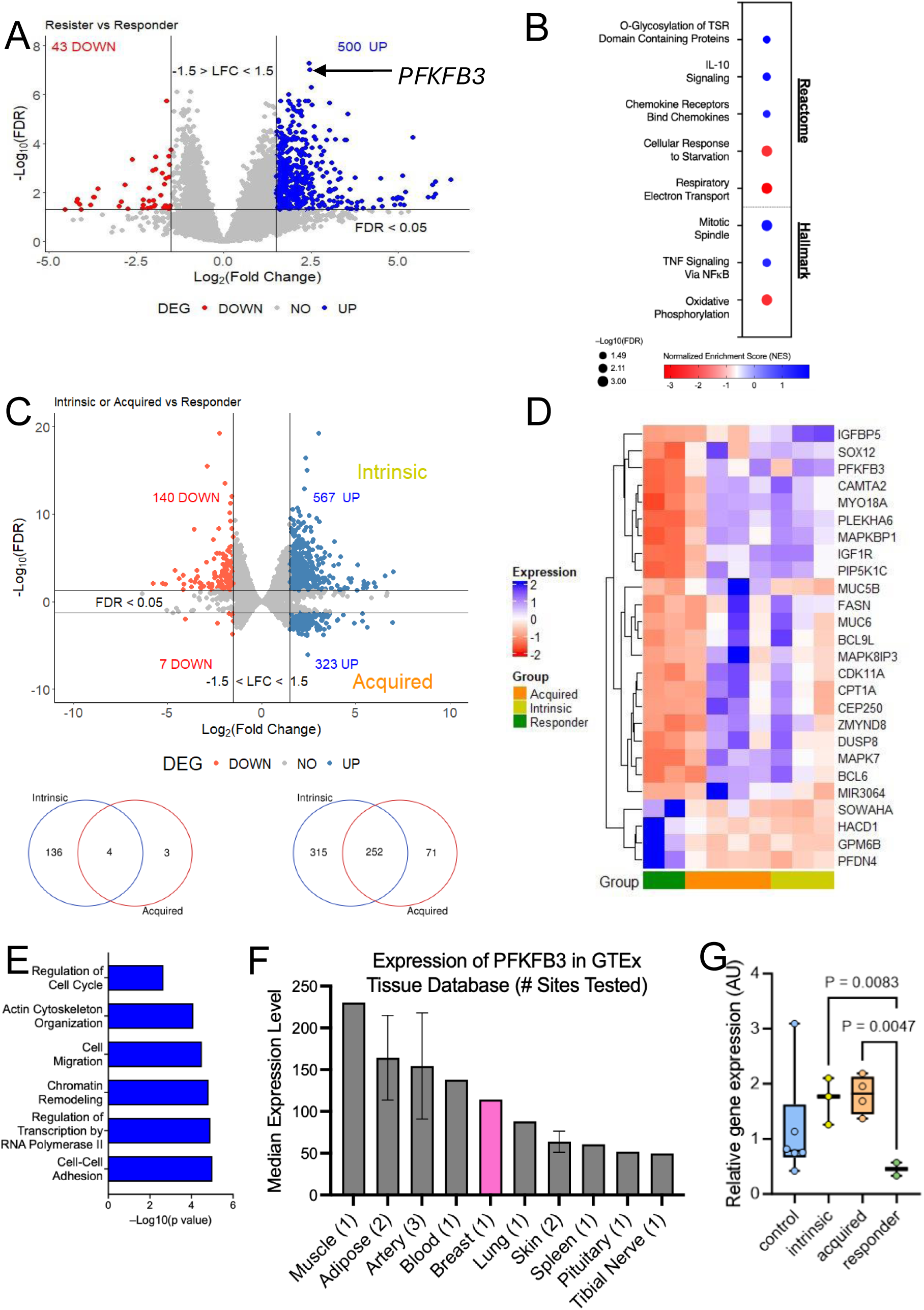

### Tumor-derived HER2+ breast cancer cells are resistant to trastuzumab and rely on HER2 signaling

To confirm the resistant nature of the tumors, we serially transplanted three trastuzumab-resistant tumors (M2 tumors T4, T6, and T7) that failed to respond to trastuzumab treatment (Fig. 1B) into treatment-naïve nude mice (Fig. 4A). Trastuzumab treatment was resumed when the tumors reached a volume of 200 mm³ (Fig. 4B). All M3 tumors derived from these xenografts remained unresponsive to trastuzumab, showing tumor growth rates and HER2 signaling comparable to those of the control tumors (Fig. 4B; Suppl. Fig. S2 A-D).

**Figure 4.**
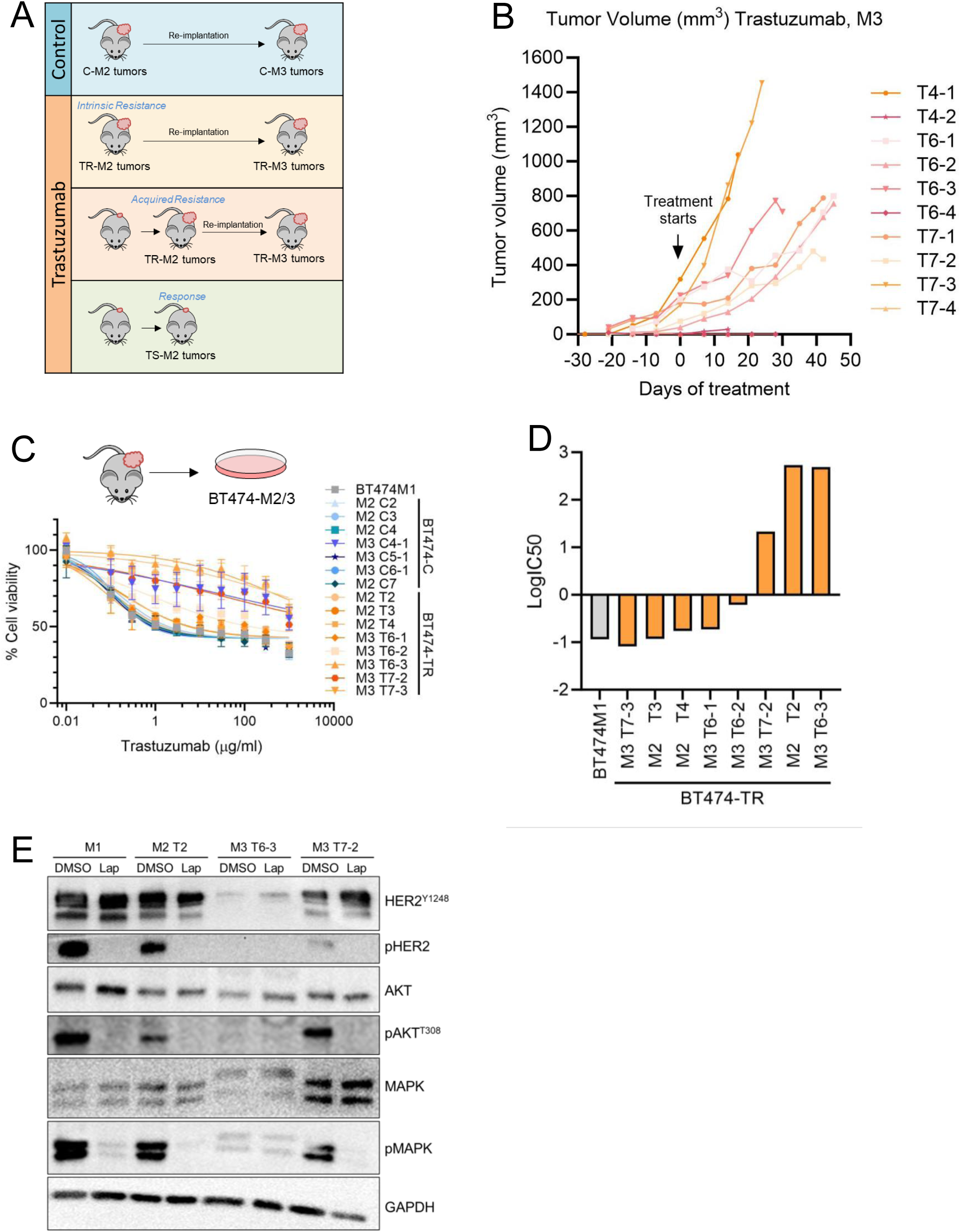

To further investigate the role of PFKFB3 as a mechanism of trastuzumab resistance, tumors were excised and cultured *ex vivo* to generate the M2 and M3 sublines. These tumor-derived cell lines, named M2-T# and M3-T# (corresponding to M2 and M3 tumors treated with trastuzumab plus tumor number), were characterized for their sensitivity to trastuzumab using cell viability assays (Fig.4 C,D). The tumor-derived cell lines exhibited varying levels of trastuzumab resistance, with IC50 ranging from 0.08 to 500 μg/ml. An IC50 higher than 1μg/ml for trastuzumab after 6-day treatment was established to define resistance in our model. This concentration was chosen based on two criteria: first, it was the lowest concentration that achieved maximum growth inhibition in parental BT474-M1 and untreated tumor-derived cell lines (Fig. 4C); second, the viability observed for the IgG control-treated cells was unaltered until high concentrations (1 mg/ml IgG) were used (Suppl. Fig. 2E).

Among the tumor-derived cell lines, three retained resistance to trastuzumab in cell culture (Fig. 4D). Importantly, resistance in the M3 T6-3 cell line could be due to downregulation of HER2. However, the other two tumor-derived resistant cell lines (M2 T2 and M3 T7-2) expressed similar HER2 levels compared to BT474-M1 cells (Fig. 4E). The growth and survival of these two cell lines remained driven by HER2, as evidenced by the inhibition of PI3K and MAPK signaling pathways upon treatment with lapatinib (Fig. 4E).

### Silencing PFKFB3 re-sensitizes trastuzumab-resistant tumor-derived HER2+ breast cancer cells

Of the eight tumor-derived cell lines, four exhibited the highest levels of PFKFB3 (Fig. 5A) and the highest resistance to trastuzumab, as indicated by their logIC50 values (Fig. 5B). These findings suggest that PFKFB3 plays a role in modulating the response of HER2+ breast cancer cells to trastuzumab.

**Figure 5.**
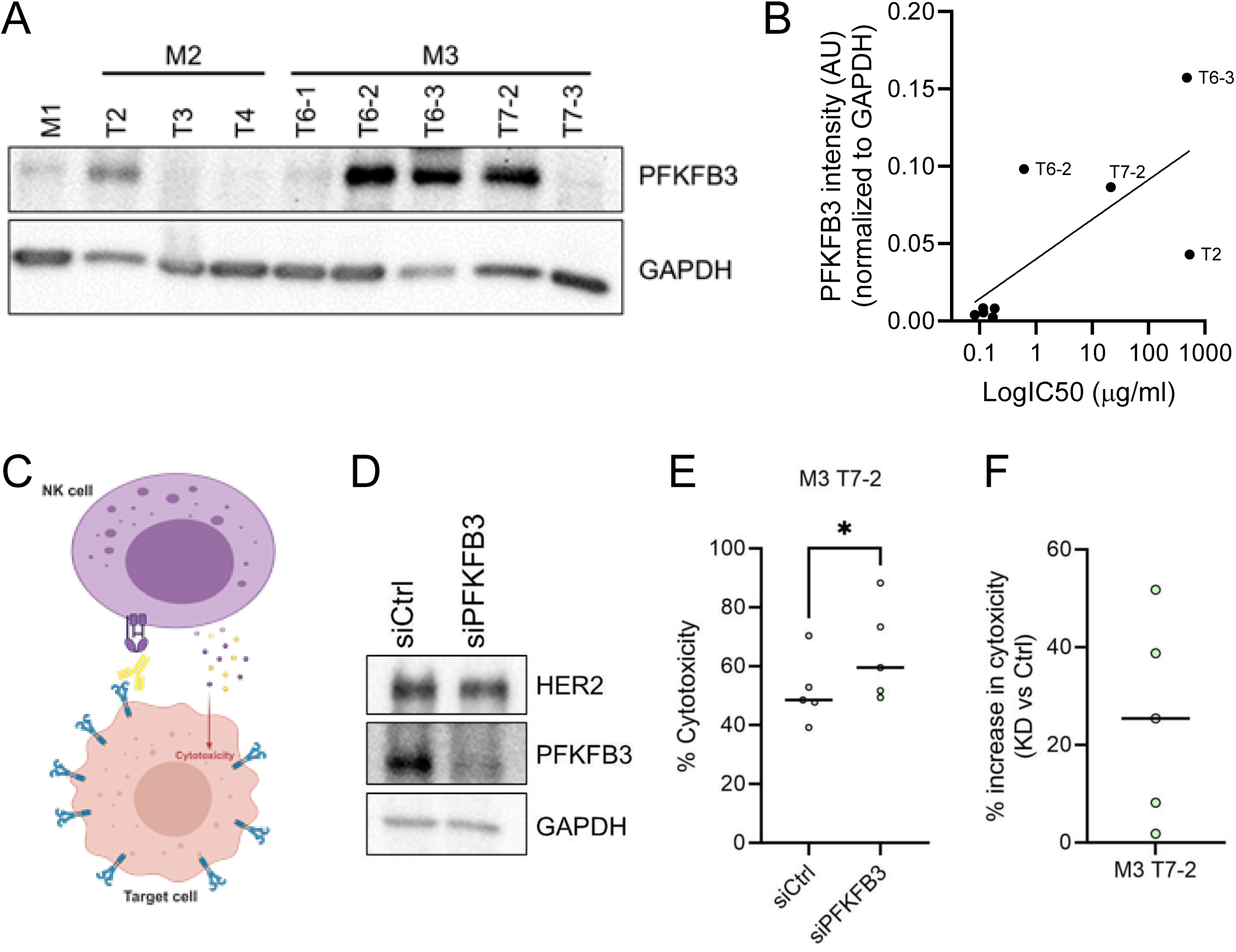

Based on these results, we selected the BT474-M3 T7-2 cell line, which maintains trastuzumab resistance and expresses high levels of *PFKFB3* for further analysis.

To evaluate the impact of reducing PFKFB3 levels on trastuzumab sensitivity, we utilized a cancer-immune co-culture system incorporating NK cells, the primary mediators of antibody-dependent cellular cytotoxicity (ADCC). Using this system (previously established by our group [25])(Fig. 5C), we knocked down *PFKFB3* in the selected tumor-derived BT474-M3 T7-2 cell line and assessed NK cell-mediated cytotoxicity against breast cancer cells. Our analysis demonstrated a significant increase in ADCC upon *PFKFB3* knockdown (Fig. 5D,E), resulting in a more efficient elimination of cancer cells (Fig. 5F).

### Silencing PFKFB3 re-sensitizes established HER2+ breast cancer cells to trastuzumab and alters their metabolism

To further validate the role of PFKFB3 in trastuzumab resistance, we assessed its expression in a broad panel of established HER2+ breast cancer cell lines (Fig. 6A,B). Among these, we selected the PFKFB3-expressing cell lines HCC1419 (from ductal carcinoma TNM stage IIIA) and HCC1569 (from metaplastic carcinoma with a germline mutation in *FHIT*) for detailed analysis. We knocked down *PFKFB3* in these two cell lines (Fig. 6C) and co-cultured them with NK cells for 24 h and 6 h, respectively. The results demonstrated a substantial increase in cell death upon *PFKFB3* knockdown in both HER2+ breast cancer cell lines, with an increase in cytotoxicity of 50% in HCC1419 cells and 40% in HCC1569 cells (Fig. 6D,E).

**Figure 6.**
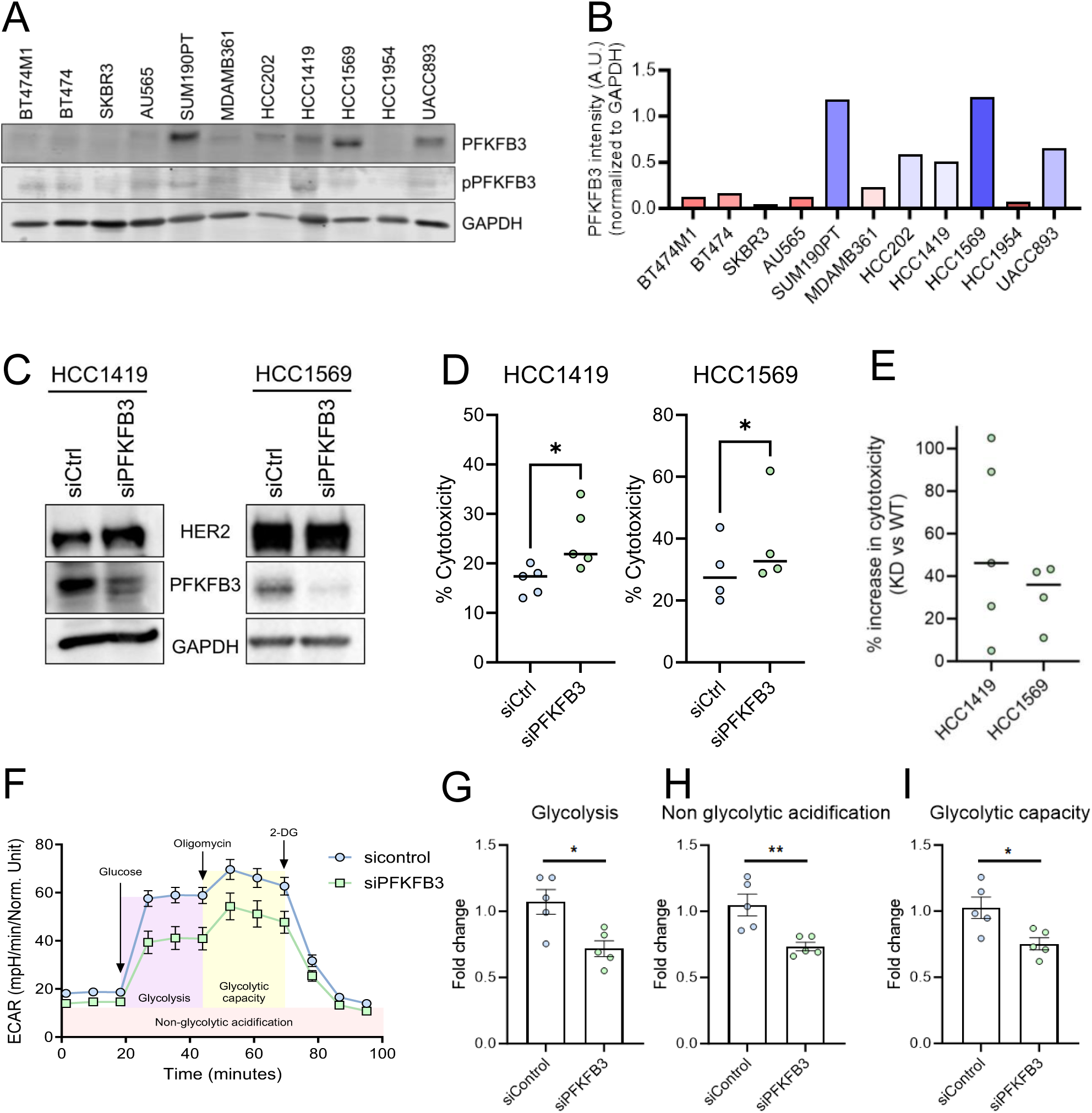

Because glycolysis was upregulated in trastuzumab-treated tumors (Fig. 2C) with *PFKFB3* upregulated in resistant tumors (Fig. 3D), we assessed the glycolytic function (ECAR) of *PFKFB3* knockdown HCC1419 cells using the glycolysis stress test by Seahorse (Fig. 6F). Knockdown of *PFKFB3* resulted in lower glycolysis, glycolytic capacity, and non-glycolytic acidification compared to control cells (Fig. 6G-I). These findings suggest that PFKFB3 sustains a glycolytic state and plays a role in trastuzumab resistance, thereby limiting NK cell-mediated cytotoxicity.

### Validation of PFKFB3 Upregulation in Clinical Cohorts

To explore the clinical relevance of PFKFB3 upregulation on trastuzumab resistance, we explored publicly available transcriptomic datasets from patients with HER2+ breast cancer. First, to investigate its prognostic relevance, using RNA-seq expression data from 379 HER2+ breast cancer patients, we observed a significant correlation between high *PFKFB3* expression and lower overall survival (OS) (Fig. 7A). This prompted us to evaluate gene chip datasets comprising 882 and 420 HER2+ patients for relapse-free survival (RFS) and OS, respectively. While OS was not statistically significant in these datasets (Supp. Fig. 3A), *PFKFB3* expression emerged as a strong predictor of RFS (Fig. 7B).

**Figure 7.**
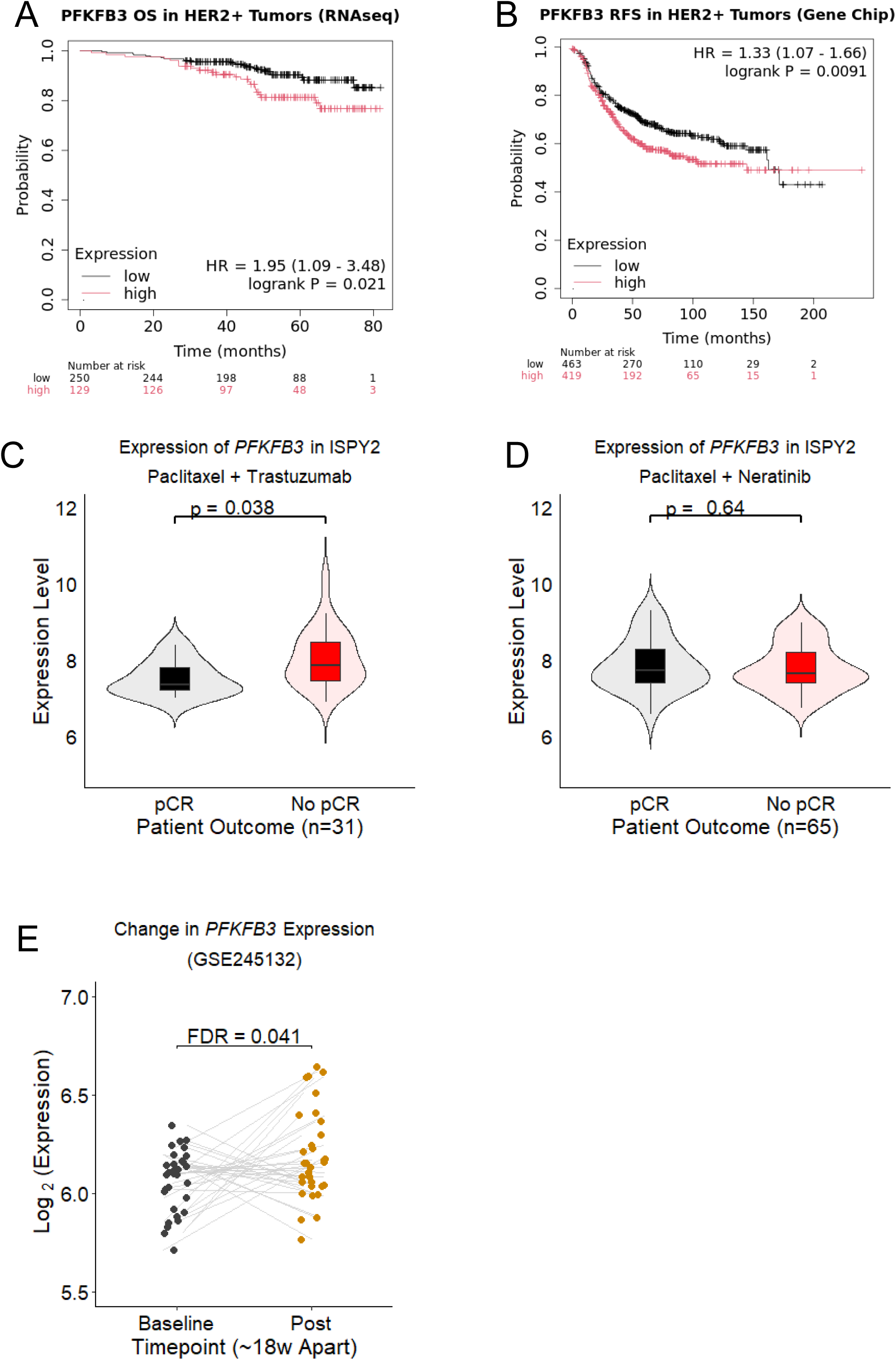

Next, we investigated the expression of *PFKFB3* in pre-treatment samples from HER2+ patients who received trastuzumab in the ISPY2 clinical trial [35]. We observed that patients achieving pathological complete response (pCR) (n=23) had lower *PFKFB3* levels than those who did not achieve pCR (n=8) in the “Trastuzumab + Paclitaxel” arm (Fig. 7C). This effect was not observed in the other trastuzumab-containing arms containing additional agents (Supp. Fig. 3B-D) nor in the arm containing the HER2-targeting tyrosine kinase inhibitor neratinib (Fig. 7D). This suggests that *PFKFB3* upregulation is a specific mechanism underlying intrinsic resistance to trastuzumab.

In two separate studies on patients after short-term treatment with trastuzumab [36, 37], no difference in *PFKFB3* expression was observed in post-treatment biopsies after 14 days in 50 patients (Supp. Fig. 3E), and after 21 days in 17 patients (Supp. Fig. 3F). However, in a study of 32 patients who did not achieve pCR after six cycles of trastuzumab plus chemotherapy over 18 weeks [38], post-treatment tumors had higher expression of *PFKFB3* than baseline biopsies (Fig. 7E), supporting that *PFKFB3* upregulation is also a specific mechanism of acquired trastuzumab resistance.

## Discussion

The mechanisms of therapeutic resistance often leverage lineage plasticity, allowing tumor cells to undergo cell state transitions that enable continued proliferation despite therapeutic pressure. Understanding these mechanisms is crucial for the development of more effective personalized treatments. In HER2+ breast cancer, various mechanisms of resistance to anti-HER2 therapies have been extensively reviewed in recent years [11, 12, 14, 15, 19, 21, 24, 41, 42].

In this study, we modelled the development of trastuzumab resistance using a cell line-derived xenograft model that yielded isogenic tumors with varying sensitivity levels, including intrinsic and acquired resistance. The utility of our xenograft model is highlighted by the identification of known resistance mechanisms among the upregulated genes in trastuzumab-resistant tumors, including *IGF1R*, *EGFR*, *FASN*, *ZMYND8*, and *CPT1A* [43–45].

Among the most significantly upregulated genes in non-responsive tumors, transcriptomic analysis identified *PFKFB3* as a new indicative gene and player contributing to trastuzumab resistance, both intrinsic and acquired. Using cancer-immune co-culture systems to examine its role in conferring resistance, we found that silencing *PFKFB3* in multiple breast cancer models re-sensitizes them to trastuzumab. Furthermore, we confirmed the clinical relevance of our findings by correlating increased *PFKFB3* expression with poorer patient outcomes in HER2+ breast cancer and lack of pCR following treatment with trastuzumab.

PFKFB3, a key regulatory enzyme in glycolysis, plays a critical role in regulating cellular metabolism [46]. Enhanced glycolysis enables cancer cells to meet increased energy demands [47] and, as a central glycolytic regulator, PFKFB3 promotes cancer cell survival, proliferation, migration, and metastasis [48–50]. Metabolic reprogramming of cancer cells is increasingly being recognized as a key feature of cancer progression and therapy resistance. While PFKFB3 primarily promotes glycolysis, other metabolic pathways, such as oxidative phosphorylation and lipid metabolism, have been implicated in cancer resistance mechanisms [51–53]. For example, a recent study demonstrated that targeting fatty acid oxidation via Cpt1a (an enzyme critical for fatty acid oxidation in the liver, breaking down fats for energy production) alongside HER2-targeting therapies combats resistance in HER2+ breast cancer preclinical models [43]. In line with this, we observed elevated *CPT1A* expression in resistant tumors. Targeting these pathways in combination with traditional therapies may provide new opportunities to overcome drug resistance and improve patient outcomes.

Our findings reveal increased PFKFB3 as a potential biomarker of resistance in HER2+ breast cancer and a targetable vulnerability to enhance trastuzumab efficacy. PFK158, a potent small-molecule inhibitor of PFKFB3, has good pharmacokinetic and pharmacodynamic properties, induces cell death, synergistically enhances chemosensitivity in endometrial cancer [54], and promotes lipophagy and chemosensitivity in gynecologic cancers [55]. This and other PFKFB3 inhibitors have shown significant growth inhibition in preclinical models of breast, lung, glioma, ovarian, gastric, and colorectal cancers [56–61].

In summary, our study identifies PFKFB3 as a key driver of trastuzumab resistance in HER2+ breast cancers. By modulating cellular metabolism, PFKFB3 enables cancer cells to adapt to therapeutic pressure and limits NK-mediated cell death. These findings pave the way for further exploration of PFKFB3 inhibitors, both as single agents and in combination with trastuzumab, to improve treatment outcomes in HER2+ breast cancer patients.

## Acknowledgments

This work in AR-S Lab was supported by a Marie Skłodowska-Curie Actions Fellowship (MSCA-IF-894191), the Ikerbasque Foundation, Victoria’s Secret Global Fund for Women’s Cancers Career Development Award, in partnership with Pelotonia & AACR (PC # 1159511); the Basque Department of Industry, Tourism, and Trade (Elkartek); the Ministerio de Ciencia e Innovación (MICINN; PID2023-150421OB-I00 and Ramón y Cajal Program); a CRIS Emerging Leader Grant from the Fundación CRIS Contra el Cáncer (PR_LE_2023-10), the VI Proyecto FERO – GHD awarded by the FERO Foundation (PFERO2024.02); and the AECC Excelencia Program PREMETACAN (EPAEC246710BIO). AC was supported by Fundación Cris Contra el Cáncer (PR_EX_2021-22), Leonardo Grant for Scientific Research and Cultural Creation (Beca Leonardo LEO23-2-10984-BBM-TRA-261), and the European Research Council (Consolidator Grant 819242). CIBERONC was co-funded with FEDER funds and funded by ISCIII. LB-B was supported by the AECC Foundation (POSTD19048BOZA).

We thank the members of the Preclinical Therapeutics Core at UCSF Helen Diller Family Comprehensive Cancer Center, Dr. John Martens for generously providing the cell lines HCC202 and UACC-893, and Dr. Maarten Fornerod for helpful discussions.

We thank Ashley Yoo (UCSF, US), Nagore Sacristan, Paula Torres (CIC bioGUNE, Spain), and David Yang Zhou and Bobbie Leijten (Erasmus MC) for their technical expertise. We also thank Franziska Linke and Wilma Teubel from Dr. Wytske van Weerden Lab at Erasmus MC for assisting with IHC staining. The expert technical advice from Saioa García Longarte, Maria Camila Salazar, and Amaia Zabala Letona of the Cancer Cell Signaling and Metabolism Lab (CIC bioGUNE) is gratefully acknowledged.

## Author Contributions

A. R. S. conceived and designed the study. A.R.S, R.V, V.S., L.B-B., and W.IJ performed the experiments and collected the data. A.R.S, R.V., S.P., V.S, W.IJ, and M.M.M. discussed the project, designed the experiments, and/or analyzed the data. A.R.S, R.V., and S.P. wrote the manuscript, which was further edited and approved by all the co-authors.

