## Supplementary figures and images for "Metabolic Alterations driven by PFKFB3 upregulation confer Resistance to Trastuzumab in HER2-Positive Breast Cancer"

### Supplementary Figure 1

#### Supplementary figure 1

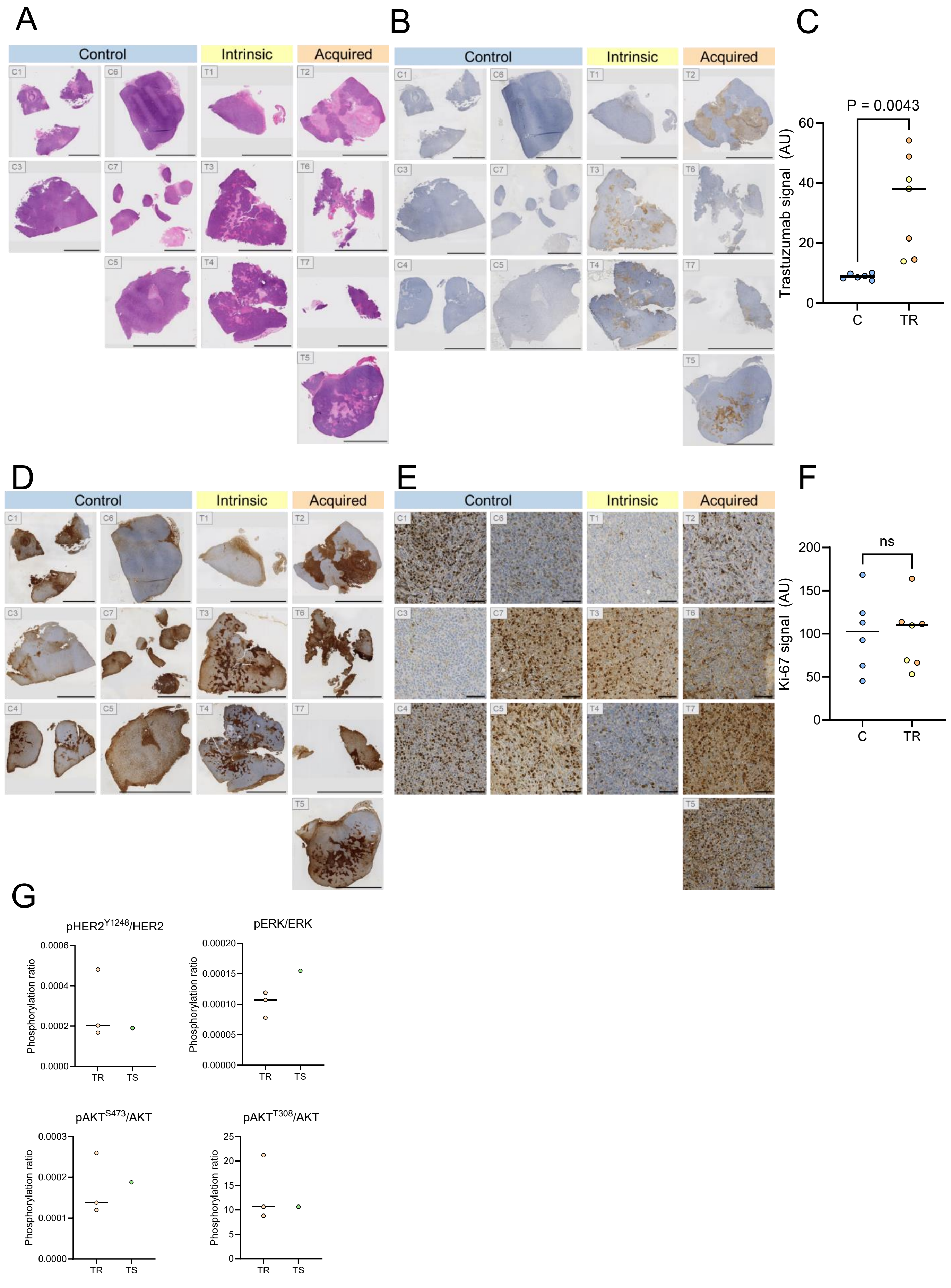

### Supplementary Figure 3

#### Supplemental Figure 3

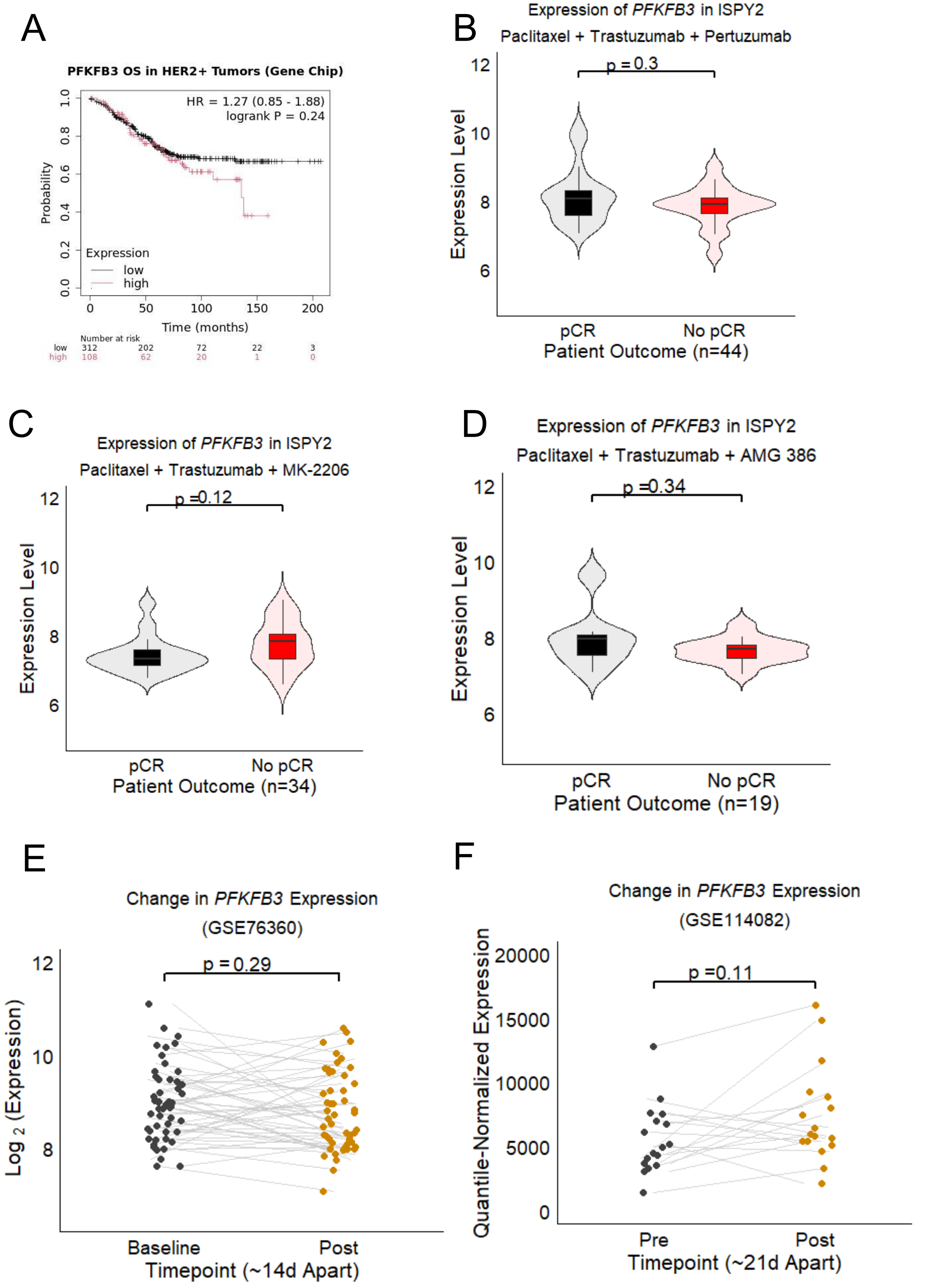
