## Supplementary Figure 2 for "Metabolic Alterations driven by PFKFB3 upregulation confer Resistance to Trastuzumab in HER2-Positive Breast Cancer"

A

| Control | | | Trastuzumab treated | | | |
|---|---|---|---|---|---|---|
| M2 Tumor | M3 derived tumors | Take rate | M2 Tumor | M3 derived tumors | Take rate | **M3 Tumors resistant to Tz treatment** |
| #C4 | #C4-1 | 1 out of 2 (50%) | #T4 | #T4-1 | 1 out of 2 (50%) | 1 out of 1 (100%) |
| #C5 | #C5-1 #C5-2 | 2 out of 3 (67%) | #T6 | #T6-1 #T6-2 #T6-3 | 3 out of 4 (75%) | 3 out of 3 (100%) |
| #C6 | #C6-1 #C6-2 | 2 out of 6 (33%) | #T7 | #T7-1 #T7-2 #T7-3 | 3 out of 4 (75%) | 3 out of 3 (100%) |
| | | 5 out of 11 (45%) | | | 7 out of 10 (70%) | **7 out of 7 (100%)** |

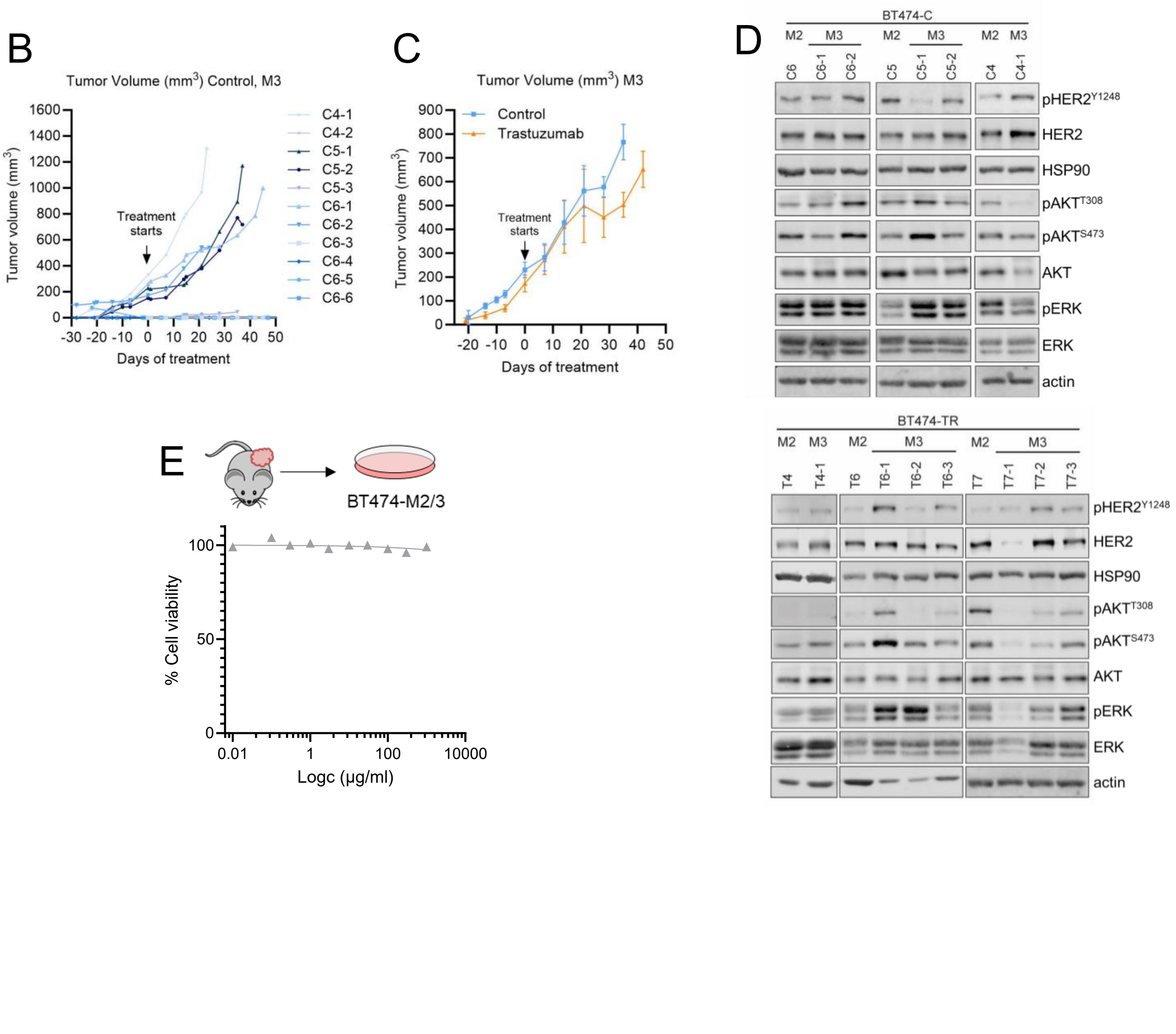
